# A Dimerization-dependent Allosteric Activation for Receptor-like Kinase in Plants

**DOI:** 10.1101/2024.01.06.574453

**Authors:** Jia Chen, Dan Cai, Yunxuan Zhang, Lin Chen, Feng Yu

**Affiliations:** State Key Laboratory of Chemo/Biosensing and Chemometrics, College of Biology, and Hunan Key Laboratory of Plant Functional Genomi-cs and Developmental Regulation, Hunan University, Changsha 410082, P.R. China

**Keywords:** Receptor-like kinase FER, kinase activity, allosteric regulation

## Abstract

Receptor-like kinases (RLKs) are essential in plants and phosphorylation is a critical step for their function. Interestingly, RLKs have many non-catalytic kinases/ pseudokinases and the biochemical basis for these pseudokinases remains unclear. FERONIA (FER) is an RLK with kinase activity, but the necessity of its kinase activity for genetic functions has been debated. Here, we uncovered that the kinase-deficient variant FER^K565R^ can activate kinase activity in FER and its homologous through homo/heterodimerization-dependent allosteric activation. We further showed that reactive oxygen species (ROS) significantly promote the dimerization of FER family members. Next, we revealed that mutating the FER P740 within the αG-αH loop reduces FER dimerization and disrupts its allosteric activation, thus attenuating FER’s transphosphorylation for its substrate. This disruption in allosteric activation abolishes the genetic function of FER^K565R^, impacting ROS production and ABA-mediated stomatal movements. Additionally, we found that MEDOS1 (MDS1), a member of the FER family, is incapable of catalyzing phosphotransfer, but can boost the kinase activity of FER and HERK1 through allosteric activation. These findings settle the debate on FER’s inactivated forms, and reveal a new mechanism for allosteric activation of RLKs via redox signaling, enhancing our understanding of pseudokinases in plants.

**One-sentence summary:** FER activates kinase activity of homologous family members through allosteric activation.

## INTRODUCTION

Receptor-like kinases (RLKs) are the largest gene family in the plant kingdom and are crucial for sensing environmental cues (1, 2). Unlike animals, where G protein-coupled receptors (GPCRs) are the most prominent membrane receptor family, RLKs dominate as the largest membrane receptors in plants. Arabidopsis, for example, has been identified over 600 RLKs, accounting for 2.5% of its protein-encoding genes (3). Most RLKs consist of three domains: an extracellular ligand-binding domain (ECD), a transmembrane (TM) helix-containing domain, and a cytoplasmic kinase domain (CD) (4). The general working model suggests that RLKs recognize and bind ligand molecules through their ECDs, leading to phosphorylation of the CD. This activation facilitates the recruitment of specific intracellular substrates, initiating downstream signaling (1, 3). Extensive research has been conducted on the mechanisms of ligand recognition by various ECDs (5, 6). However, despite progress in understanding ECD-ligand and CD-substrate interactions, the specific activation process and subsequent substrate phosphorylation events in the early stages of RLK signaling remain largely unknown. This question will become more complex when we consider the many non-catalytic kinases/pseudokinase RLKs. Pseudokinase RLKs predicted to lack phosphoryl group transfer activity due to degradation of one or more highly conserved motifs that are crucial for substrate orientation and/or catalysis (7), have posed a mystery regarding their biological role in cellular transduction.

Although pseudokinases lack catalytic residues, certain members exhibit structural similarity to activate kinases and perform essential genetic role (7). In the plants, a specific subset of pseudokinases known as non-arginine-aspartate (non-RD) kinases which lack the arginine residue in the HRD motif (8). Some non-RD kinases promote antibacterial immunity independently of their phosphotransfer activity (9). For instance, the pathogen recognition receptor BIR2 in plants possesses an intracellular pseudokinase domain that undergoes phosphorylation by the associated receptor BAK1 (10). This phosphorylation event triggers a conformational change, altering the interaction properties of BIR2 and driving signaling (10). Another plant pseudokinase BSK8 exhibits unusual conformational arrangements of nucleotide phosphate groups and catalytic key motifs, which regulates brassinosteroid signal transduction through the binding and activation of the BRI1 suppressor 1 phosphatase BSU1 (11, 12). Similarly, the tomato atypical receptor kinase 1 (TARK1), another plant pseudokinase, exhibits similar characteristics to BSK8 (13). These findings emphasize pseudokinases have essential genetic role despite their lack of enzymatic activity via a still unknown biochemical mechanism.

FER is a member of the *Cr*RLK1L family of RLKs in Arabidopsis, consisting of 17 members. Extensive genetic analyses have demonstrated FER’s crucial role in plant immunity, growth, and reproduction (14, 15). Previous studies have revealed that FER interacts with various proteins through its cytoplasmic domain, such as GEF1/4/10, ATL6, eIF4E, RIPK, ABI2, EBP1, MYC2 and GRP7, to regulate different aspects of plant physiology (16–22). Although FER has been shown to have numerous substrates, the exact role of its kinase activity in signal transduction remains debatable. Studies with the kinase-deficient mutant FER^K565R^ have revealed that the kinase domain is necessary for pollen tube reception, but its activity is not required (23). Similarly, the restoration of defective ovule fertilization in the *FER^K565R^/fer-4* mutant indicates a partial independence of kinase activity for certain responses (24). Additional research has shown dose-dependent effects of FER^K565R^ on rosette morphology and stomatal movements (25). Recent study has revealed that the kinase domain of FER (FER-KD) adopts an activated conformation without relying on phosphorylation and undergoes trans-autophosphorylation, providing insights into the activation and regulation of FER and setting a paradigm for RLKs signaling initiation (26). Moreover, the structural analysis of FER-KD^K565R^ reveals its adoption of an active conformation, closely resembling that FER-KD, indicated that FER-KD^K565R^ may maintain its capability to partially recruit or activate specific downstream substrates for signal transduction, similar to some pseudokinases (26). Nevertheless, the genetic and biochemical implications behind the active conformation of both FER-KD and FER-KD^K565R^ remain unclear at present.

In this study, following our previous work (26), we found the kinase-deficient variant FER^K565R^ and MDS1 activated the kinase activity of FER homologous members through allosteric regulation. Furthermore, mutation of the FER dimerization site P740 within αG-αH loop resulted in a significant reduction in FER’s transphosphorylation ability and exhibited a FER partial loss-of-function phenotype. These findings uncover a novel biochemical regulatory mechanism of RLKs and resolve the longstanding debate regarding the capacity of inactivated forms of FER to still exert partial genetic function output.

## RESULTS

### FER^K565R^ and MDS1 enhance the kinase activity of FER and HERK1 through allosteric activation

Based on previous genetic results, the kinase-deficient mutant FER^K565R^ exhibits dose-dependent effects on the rosette morphology and RALF1-mediated stomatal movements (25). To validate whether FER^K565R^ enhances FER’s kinase activity, we used the previously reported FER phosphorylation substrate glycine-rich RNA-binding protein 7 (GRP7) (21) as the substrate to verify the allosteric activation ability of FER^K565R^. Based on *in vitro* kinase assays, FER-KD^K565R^ facilitated transphosphorylation of FER-CD in a concentration-dependent manner (Figure 1A). With the concentration of FER-KD^K565R^ increasing, the phosphorylation of GRP7 proportionally increased (Figure 1A). This finding suggested that despite lacking intrinsic kinase activity, FER-KD^K565R^ is able to activate the transphosphorylation capacity of FER-CD towards GRP7 through a still unknown mechanism. To further confirm this mechanism, we performed titration experiments by adding increasing amounts of FER^K565R^ into *Ubi::FER-FLAG* protoplasts. We induced FER kinase activity by externally introducing RALF1 peptide and monitored the phosphorylation of GRP7. The result showed that elevated levels of FER^K565R^ corresponded to an increased phosphorylation of GRP7 (Figure 1B).

**Fig. 1.**
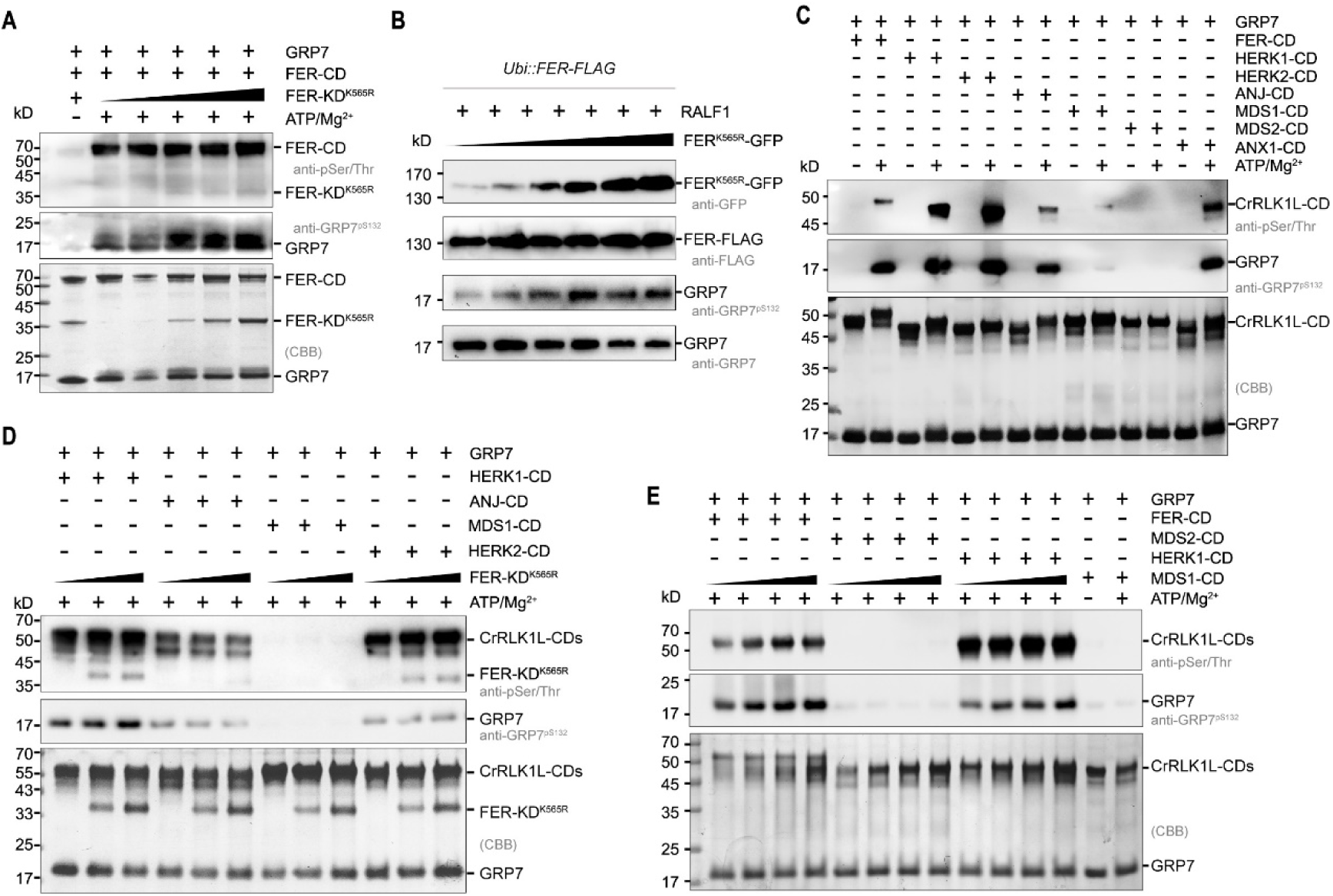
FER^K565R^ and MDS1 enhance the kinase activity of FER and HERK1. (**A**) Concentration gradient analysis of FER-CD^K565R^ trans-phosphorylates FER-CD and GRP7 substrate. Samples were incubated with 1 mM ATP and 10 mM Mg^2+^ at 25 ℃ for 30 min. (**B**) Protoplasts were prepared from 7-day-old *Ubi::FER-FLAG* seedlings. Concentration gradient of FER^K565R^-GFP (0, 2, 5, 10, 15, 20 or 40 μg) DNA plasmids were transfected into *Ubi::FER-FLAG* protoplasts. The transfected protoplasts were treated with 1 μM RALF1 for 15 min. Total protein of the protoplasts were extracted and analyzed by SDS‒PAGE and immunoblotting with anti-FLAG, anti-GRP7, anti-GRP7^pS132^ and anti-GFP antibodies. (**C**) Western blot analysis of autophosphorylation and transphosphorylation of FER-CD, HERK1-CD, HERK2-CD, ANJ-CD, MDS1-CD, MDS2-CD and ANX1-CD. Samples were incubated with 1 mM ATP and 10 mM Mg^2+^ at 25°C for 30 min. (**D**) Concentration gradient analysis of FER-KD^K565R^ transphosphorylases HERK1-CD, HERK2-CD, ANJ-CD, and MDS1-CD. Samples were incubated with 1 mM ATP and 10 mM Mg^2+^ at 25 ℃ for 30 min. (**E**) Concentration gradient analysis of MDS1-CD transphosphorylases FER-CD, HERK1-CD, and MDS2-CD. Samples were incubated with 1 mM ATP and 10 mM Mg^2+^ at 25 ℃ for 30 min.

To further investigate the activating effect of FER^K565R^ on the kinase activity of FER homologs, we purified HERK1-CD, HERK2-CD, ANJ-CD, ANX1-CD, MDS1-CD and MDS2-CD (Figure S1A). *In vitro* kinase assays demonstrated that HERK1-CD, HERK2-CD, ANJ-CD and ANX1-CD displayed autophosphorylation activity and facilitated the phosphorylation of GRP7 (Figure 1C). However, MDS1-CD and MDS2-CD lacked obvious autophosphorylation activity and was unable to trans-phosphorylate GRP7 (Figure 1C). Increasing concentrations of FER-KD^K565R^ resulted in the augmentation of transphosphorylation of HERK1-CD (Figure 1D). Nevertheless, FER-KD^K565R^ failed to enhance the kinase activity of ANJ-CD or induce kinase activity of MDS1-CD (Figure 1D). *In vitro* kinase assays showed that FER homologous family members, MDS1 and MDS2, do not exhibit autophosphorylation activity. The question of whether they possess allosteric activation regulatory capabilities similar to FER-KD^K565R^ was raised. The findings indicated that, consistent with FER-KD^K565R^, MDS1 was capable of concentration-dependent enhancement of the kinase activity of FER and HERK1 (Figure 1D), thereby confirming MDS1’s ability to allosterically activate homologous family members.

### FER and its homologs undergo both homologous and heterologous dimerization in responding to ROS signaling

The dimerization is a critical prerequisite and foundation for allosteric regulation. To investigate the dimerization of FER and its homologs, we performed chemical crosslinking on FER-CD and its homologs in the presence of an amine-to-amine crosslinker, disuccinimidyl suberate (DSS, A reagent commonly used for the study of protein dimerization). We observed that FER-CD, MDS1-CD, and ANX1-CD exhibit strong dimerization and oligomerization, with FER-CD^K565R^ showing a higher degree of dimerization compared to FER-CD (Figure 2A). In contrast, homologous dimerization of HERK1-CD, ANJ-CD and MDS2-CD appeared relatively weaker (Figure 2A). Further heterologous dimerization experiments indicated that FER-CD^K565R^ and MDS1-CD can form heterologous dimers with homologous family members without the need for DSS induction (Figure 2B), implying that the concentration-dependent promotion of kinase activity for FER and HERK1 is mediated by intermolecular heterologous dimerization. This suggests that FER may regulate its kinase activity through an intermolecular dimerization mechanism.

**Fig. 2.**
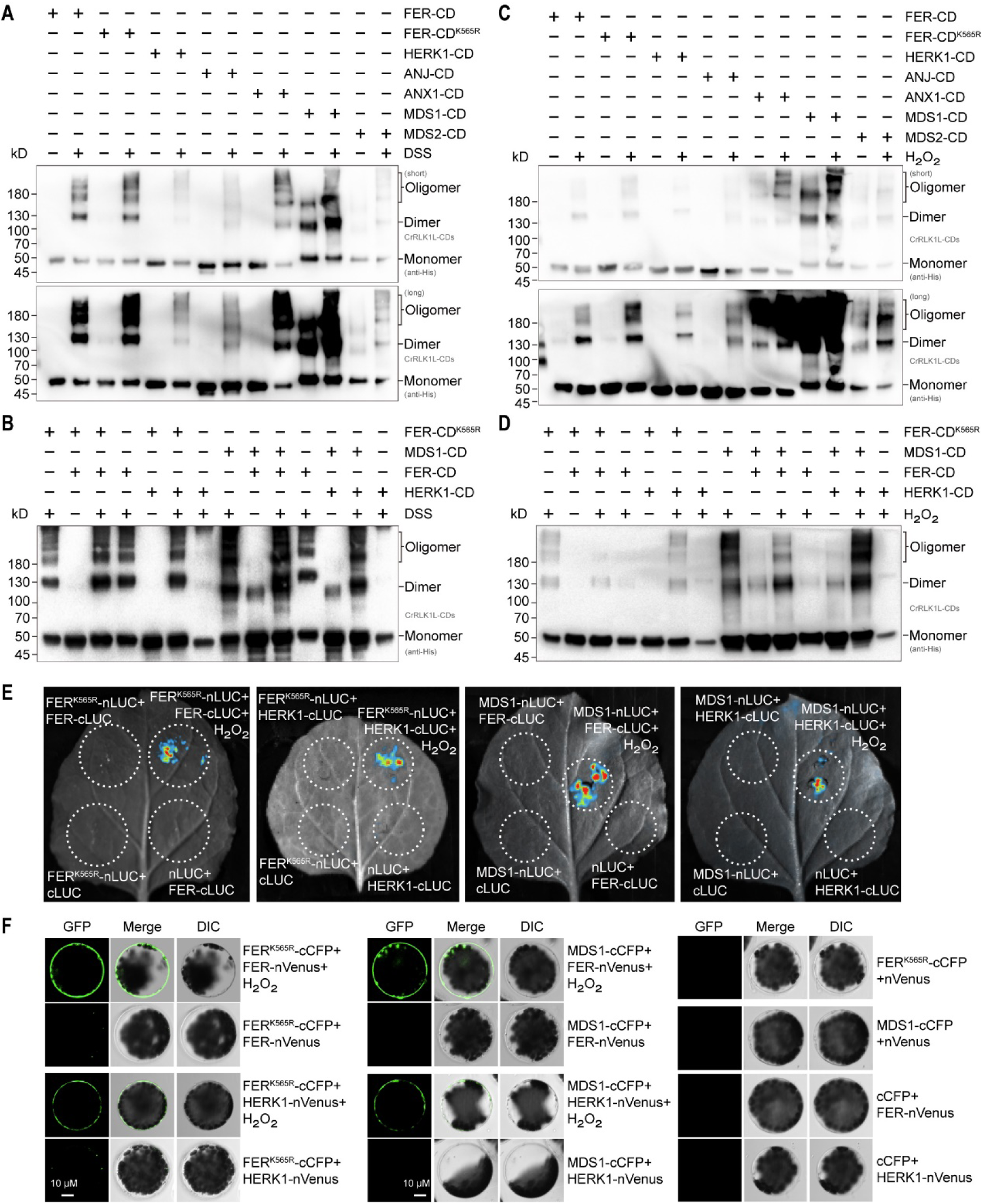
Homologous and heterologous dimerization in FER and its homologs following oxidation. (**A**) Homologous dimerization analysis of FER-CD and its homologous family incubated with DSS for 30 min. The monomer and dimer/oligomer were detected using α-anti-His. (**B**) Heterologous dimerization analysis of FER-CD^K565R^ or MDS1 with its homologous family FER-CD and HERK1-CD incubated with 100 μM DSS for 30 min. The monomer and dimer/oligomer were detected using α-anti-His. (**C**) Homologous dimerization analysis of FER-CD and its homologous family incubated with 0.5 mM H_2_O_2_ for 30 min. The monomer and dimer/oligomer were detected using α-anti-His. (**D**) Heterologous dimerization analysis of FER-CD^K565R^ or MDS1 with its homologous family FER-CD and HERK1-CD incubated with 0.5 mM H_2_O_2_ for 30 min. The monomer and dimer/oligomer were detected using α-anti-His. (**E**) Dimerization in FER and its homologous family following oxidation in luciferase complementation assay (LCA) assay in *N. benthamiana*. Negative controls (FER^K565R^-nLUC + cLUC, nLUC + FER-cLUC, nLUC + HERK1-Cluc, and MDS1-nLUC + cLUC) are shown. (**F**) Dimerization in FER and its homologous family following oxidation in the bimolecular fluorescence complementation (BiFC) assay in Arabidopsis protoplasts. Negative controls (FER^K565R^-cCFP + nVenus, MDS1-cCFP + nVenus, cCFP + FER-nVenus and cCFP + HERK1-nVenus) are shown. Green fluorescent protein (GFP), merged signal of GFP and differential interference contrast (DIC) are shown, bar = 10 μm.

Previous studies have shown that reactive oxygen species (ROS) signaling is the key factor for FER and its homologs to perform their genetic output (27–30). ROS are reactive forms of molecular oxygen including the hydroxyl radical, superoxide, hydrogen peroxide (H_2_O_2_) and singlet oxygen (31). Therefore, we aimed to verify whether FER and its homologs are subjected to oxidative dimerization regulation. FER and its homologs contain varying numbers of cysteine residues, and there are significant differences in the distribution of cysteine residues (Figure S2). We found that after treatment with H_2_O_2_, FER-CD, FER-CD^K565R^, ANX1-CD, and MDS1-CD exhibited significant dimerization (Figure 2C), while the dimerization of HERK1-CD, ANJ-CD, and MDS2-CD was less pronounced (Figure 2C), similar to the dimerization results observed after DSS treatment. Further experiments revealed that FER-CD^K565R^ and MDS1-CD can form heterologous dimers with FER-CD and HERK1-CD after treated with H_2_O_2_ (Figure 2D), suggesting that ROS signaling potentially plays a critical role in the allosteric regulation and kinase activity of FER and its homologs. To further validate ROS-induced FER and its homologs dimerization *in vivo*, we performed bimolecular fluorescence complementation (BiFC) and luciferase complementation assay (LCA), and found that H_2_O_2_ indeed promoted homologous dimerization of FER and heterologous dimerization of FER with HERK1 (Figure 2E and F).

### FER face-to-face interface residues are crucial for its dimerization

Based on our previous work analysis of the FER-KD crystal structure (26), we observed two distinct symmetric dimerization conformations of FER-KD: face-to-face and back-to-back (Figure 3A and B). The face-to-face dimerization interface consists of 6 critical amino acid residues, namely R711, R712, P740, L742, E751 and D792 (Figure 3A). Similarly, the back-to-back dimerization interface involves seven key amino acid residues, including R589, R591, Y648, G652, A653, D802 and W805 (Figure 3B). To investigate the actual dimerization of FER, site-directed mutagenesis was employed to introduce specific alanine (A) substitutions at the face-to-face dimer interface residues (R711A, R712A, P740A, L742A, E751A and D792A) (Figure S1B). Sites-directed mutagenesis was employed to introduce alanine (A) substitutions (R589A, R591A, D802A and W805A), glutamine (N) substitution (G652N) and glutamic acid (D) substitution (A653D) at the back-to-back dimer interface residues (Figure S1B). We performed chemical crosslinking on FER-CD and face-to-face dimer interface mutants in the presence of DSS, found that the degree of P740A dimerization is significantly reduced (Figure 3C), while mutations of R712, L742, E751 and D792 have little impact on FER dimerization (Figure 3C). Further analysis of back-to-back dimer interface mutants reveals that back-to-back dimer interface residues have minimal impact on FER dimerization (Figure 3D).

**Fig. 3.**
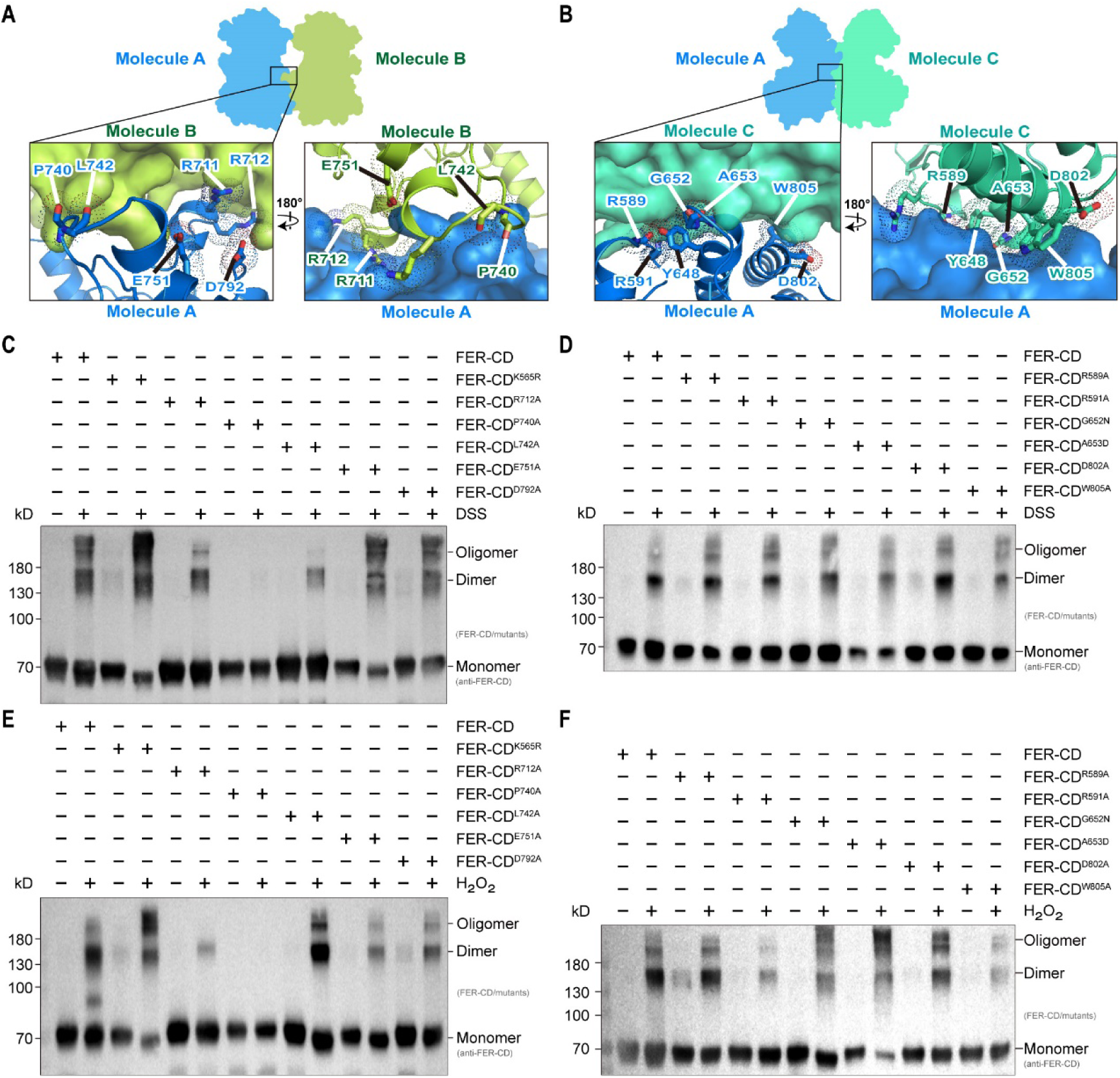
FER face-to-face interface residues are crucial for its dimerization. (**A**) Overview of the face-to-face dimer in the crystal and detailed view of the dimeric interface. Molecule B (green-yellow) is shown in surface representation in the first exploded view. Interfacial residues from molecule A (dark blue) are highlighted. In the second exploded view, molecule A is shown in surface representation, and interfacial residues from molecule B are highlighted. (**B**) Overview of the back-to-back dimer in the crystal and detailed view of the dimeric interface. Molecule C (green–cyan) is shown in surface representation in the first exploded view. Interfacial residues from molecule A are highlighted. In the second exploded view, molecule A is shown in surface representation, and interfacial residues from molecule C are highlighted. (**C**) Dimerization analysis of FER-CD face-to-face mutations and K565R incubated with DSS for 30 min. The FER-CD monomer and dimer/oligomer were detected using anti-FER-CD. (**D**) Dimerization analysis of FER-CD back-to-back mutations incubated with DSS for 30 min. The FER-CD monomer and dimer/oligomer were detected using anti-FER-CD. (**E**) Dimerization analysis of FER-CD face-to-face mutations and K565R incubated with 0.5 mM H_2_O_2_ for 30 min. The FER-CD monomer and dimer/oligomer were detected using α-anti-FER-CD. (**F**) Dimerization analysis of FER-CD back-to-back mutations incubated with 0.5 mM H_2_O_2_ for 30 min. The FER-CD monomer and dimer/oligomer were detected using anti-FER-CD.

To further validate whether mutations at the face-to-face dimerization interface sites affect the dimerization after oxidation, we conducted dimerization validation induced by H_2_O_2_. Our findings indicated that FER-CD undergone oxidation-induced dimerization (Figure 3E), while FER-CD^K565R^ experience a reduction in monomeric molecular weight after oxidation, accompanied by a noticeable increase in dimerization and oligomerization (Figure 3E). This suggests that oxidation promotes the formation of intramolecular disulfide bonds, thereby facilitating dimerization and oligomerization. Conversely, FER-CD^R712A^ and FER-CD^P740A^ exhibited decreased dimerization and oligomerization after oxidation, particularly FER-CD^P740A^, which completely lost its dimerization ability (Figure 3E). This underscores the pivotal role of the P740 in influencing FER’s face-to-face dimerization. When we analyzed the P740 site in detail, we found that P740 within the αG-αH loop, in the center of the loop implies its significance in the regulation of dimerization (Figure S3). The back-to-back dimer interface mutants had minimal impact on FER dimerization after H_2_O_2_ treatment (Figure 3F). Consequently, these findings emphasize the critical significance of residues P740 within the face-to-face interface for FER’s dimerization, suggesting that FER may regulate its kinase activity through an intermolecular dimerization mechanism.

### The dimerization of FER-CD is required for FER and HERK1 kinase activity

To validate the dimerization of FER-CD induced by H_2_O_2_ and its impact on FER kinase activity, we proceeded with *in vitro* kinase assays and found that, H_2_O_2_ promoted FER-CD’s autophosphorylation and GRP7’s transphosphorylation in a concentration-dependent manner, whereas DTT treatment inhibited these processes in a concentration-dependent manner (Figure 4A), emphasizing the importance of dimerization and allosteric regulation in FER kinase activity. We further conducted kinase activity verification on FER homologs, and found that, similar to FER-CD, H_2_O_2_ significantly enhances the kinase activity of HERK1-CD (Figure 4B). However, after DTT treatment, the kinase activity of HERK1-CD markedly decreased (Figure 4B). In contrast, the kinase activities of HERK2-CD and ANJ-CD remain insensitive to both H_2_O_2_ and DTT treatments (Figure 4B), further indicating that the dimerization of HERK1-CD is required for HERK1-CD kinase activity. To further investigate the impact of FER dimerization on FER’s kinase activity, we first analyzed the impact of residue mutations at the interface of the face-to-face dimerization on the activity of FER kinase. Based on our *in vitro* kinase assay, FER-CD exhibited robust kinase activity, while the R712A, P740A and E751A led to decreased autophosphorylation and transphosphorylation ability towards GRP7 (Figure 4C). The R711A, L742A and D792A had no significant impact on transphosphorylation of GRP7 (Figure 4C). These findings highlight the crucial importance of the residues R712A, P740A and E751A within the face-to-face dimer interface for FER’s kinase activity. In contrast, mutations of key residues within the back-to-back dimer interface (R589A, R591A, G652N, A653D, D802A, and W805) had minimal impact on FER-CD’s kinase activity (Figure S4).

**Fig. 4.**
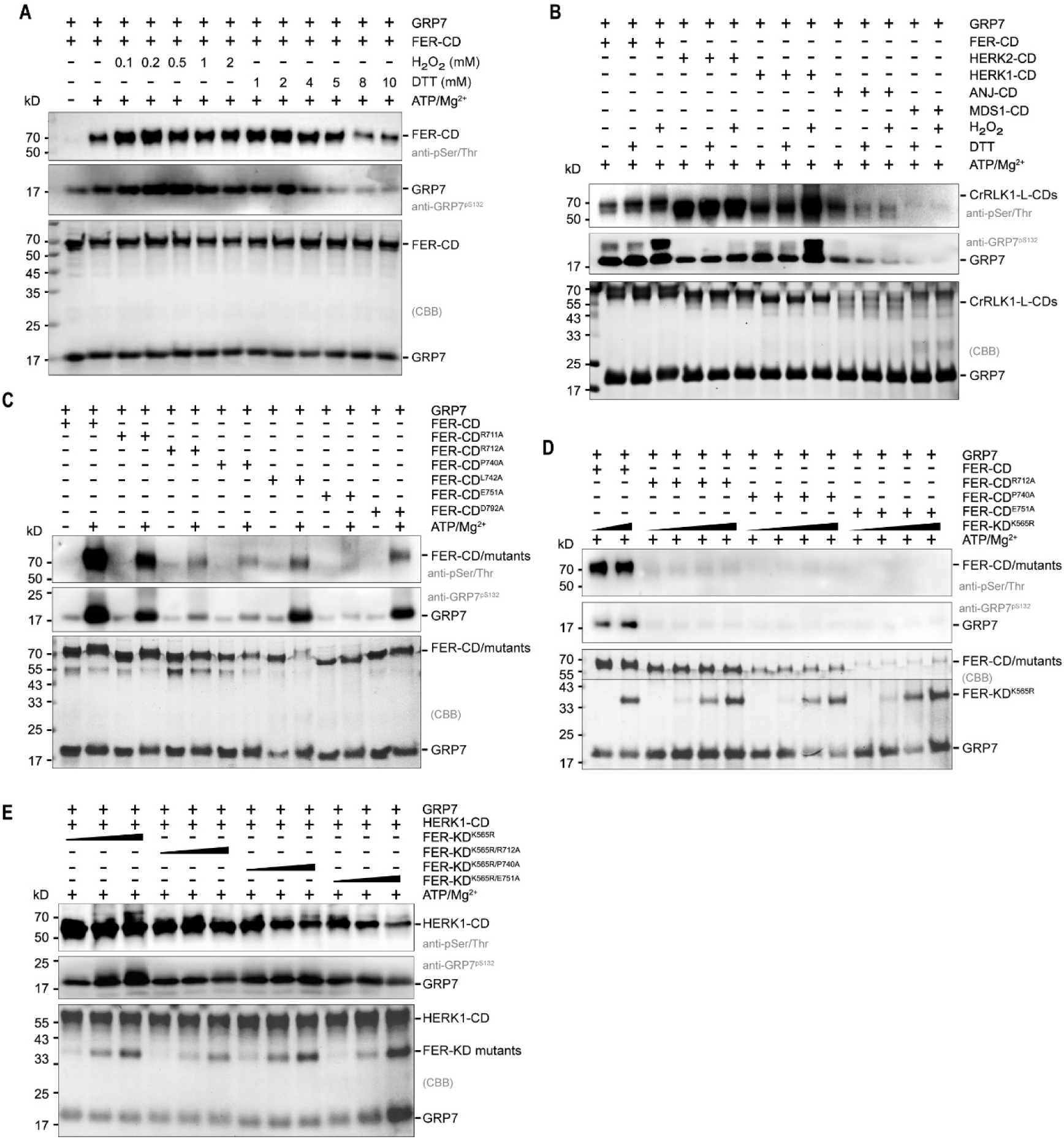
The dimerization interface sites of FER-CD are crucial for FER and HERK1 kinase activity. (**A**) Western blot analysis of autophosphorylation and transphosphorylation of FER-CD treated with different concentrations of either H_2_O_2_ or DTT. Samples were incubated with 1 mM ATP and 10 mM Mg^2+^ at 25°C for 20 min. (**B**) Western blot analysis of autophosphorylation and transphosphorylation of FER-CD, HERK1-CD, HERK2-CD, ANJ-CD or MDS1-CD treated with H_2_O_2_ or DTT. Samples were incubated with 1 mM ATP and 10 mM Mg^2+^ at 25°C for 20 min. (**C**) Western blot analysis of autophosphorylation and transphosphorylation of face-to-face dimerization interface sites of FER-CD. Samples were incubated with 1 mM ATP and 10 mM Mg^2+^ at 25°C for 30 min. (**D**) Concentration gradient analysis of FER-KD^K565R^ transphosphorylases FER-CD, FER-CD^R712A^, FER-CD^P740A^ and FER-CD^E751A^. Samples were incubated with 1 mM ATP and 10 mM Mg^2+^ at 25 ℃ for 30 min. (**E**) Concentration gradient analysis of FER-KD^K565R^, FER-KD^K565R/R712A^, FER-KD^K565R/P740A^ and FER-KD^K565R/E751A^ transphosphorylase HERK1-CD. Samples were incubated with 1 mM ATP and 10 mM Mg^2+^ at 25 ℃ for 30 min.

We further observed that FER-KD^K565R^ failed to facilitate kinase activity of FER-CD after the face-to-face dimer interface residues R712, P740 and E751 were substituted by A (Figure 4D). Thus, it is postulated that FER-CD likely utilizes face-to-face dimerization as a mechanism to facilitate substrate-specific activation. To further investigate the potential conserved role of FER’s dimerization interface in the activation of HERK1, we introduced mutations at the crucial residues of the dimerization interface (R712A, P740A and E751A) in the background of FER^K565R^. The results demonstrated that, following mutations at the key residues of the dimerization interface, FER-KD^K565R^ lost its ability to induce the activation of HERK1 (Figure 4E). Collectively, these findings strongly support the proposition that FER can activate its own kinase activity and that of the HERK1 through dimerization-mediated allosteric regulation.

### The kinase activity of dimerization site mutations of FER strongly compromised in Arabidopsis

To further investigate the role of face-to-face dimerization in activating FER’s kinase activity *in vivo*, we conducted validation experiments involving the transfection of FER-GFP, FER^K565R^-GFP, FER^R712A^-GFP, FER^P740A^-GFP, FER^L742A^-GFP, FER^E751A^-GFP and FER^D792A^-GFP constructs into the *fer-4* mutant protoplasts. FER kinase activity was induced by treatment with RALF1, and the phosphorylation of GRP7 was monitored. Notably, RALF1 significantly augmented the transphosphorylation of GRP7 by FER-GFP (Figure 5A), while FER^R712A^-GFP, FER^P740A^-GFP, and FER^E751A^-GFP abolished the pronounced phosphorylation of GRP7 induced by RALF1 (Figure 5A). To further dissect the mechanism through which FER^K565R^ activates FER’s kinase activity via face-to-face dimerization, titration experiments were performed. Increasing amounts of FER^K565R^ were incrementally introduced in conjunction with co-transfection of FER^R712A^-GFP, FER^P740A^-GFP, FER^E751A^-GFP or FER-GFP constructs into *fer-4* mutant protoplasts. We observed that elevated levels of FER^K565R^ expression correlated with an enhanced phosphorylation of GRP7 when co-transfected with FER-GFP (Figure 5B). However, increased levels of FER^K565R^ expression failed to elevate the phosphorylation of GRP7 when co-transfected with FER^R712A^-GFP, FER^P740A^-GFP or FER^E751A^-GFP (Figure 5B). To further underscore the importance of face-to-face dimerization for FER kinase activity and FER^K565R^ allosteric activation *in vivo*, we generated FER^K565R/R712A^-GFP, FER^K565R/P740A^-GFP and FER^K565R/E751A^-GFP and performed titration experiments. Our findings revealed that increased levels of FER^K565R/R712A^-GFP, FER^K565R/P740A^-GFP, or FER^K565R/E751A^-GFP expression did not enhance GRP7 phosphorylation when transfecting into *Ubi::*FER-FLAG protoplasts (Figure 5C). These experimental findings collectively suggest that the face-to-face dimer interface of FER plays a pivotal role in its allosteric activation, ultimately modulating FER’s kinase activity.

**Fig. 5.**
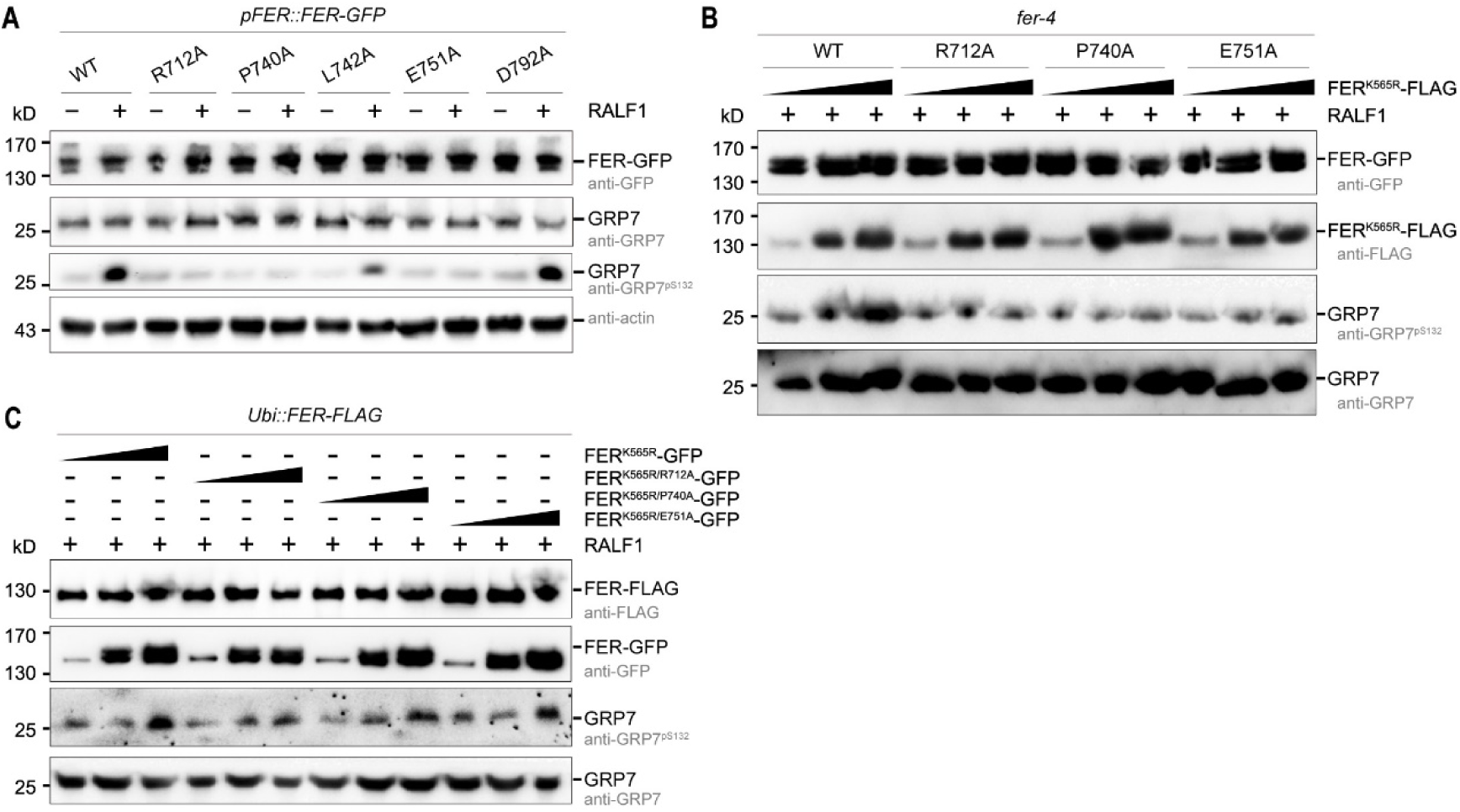
Kinase activity of the dimerization interface sites mutants are strongly compromised in Arabidopsis. (**A**) Protoplasts were prepared from 7-day-old *fer-4* seedlings. *FER-GFP*, *FER-GFP^R712A^*, *FER-GFP^P740A^*, *FER-GFP^L742A^*, *FER-GFP^E751A^* and *FER-GFP^D792A^* DNA plasmids were transfected into *fer-4* protoplasts, respectively. The transfected protoplasts were treated with 1 μM RALF1 for 15 min. Total protein of the protoplasts were extracted and analyzed by SDS‒PAGE and immunoblotting with anti-GRP7, anti-GRP7^pS132^, anti-actin and anti-GFP antibodies. (**B**) Protoplasts were prepared from 7-day-old *fer-4* seedlings. *FER-GFP*, *FER-GFP^R712A^*, *FER-GFP^P740A^* and *FER-GFP^E751A^* DNA plasmids were co-transfected with different concentrations of *FER^K565R^-FLAG* into *fer-4* protoplasts, respectively. The transfected protoplasts were treated with 1 μM RALF1 for 15 min. Total protein of the protoplasts were extracted and analyzed by SDS‒PAGE and immunoblotting with anti-FLAG, anti-GRP7, anti-GRP7^pS132^ and anti-GFP antibodies. (**C**) Protoplasts were prepared from 7-day-old *Ubi::FER-FLAG* seedlings. *FER^K565R^-GFP*, *FER-GFP^K565R/R712A^*, *FER-GFP^K565R/P740A^*, and *FER-GFP^K565R/E751A^*DNA plasmids were transfected into *Ubi::FER-FLAG* protoplasts, respectively. The transfected protoplasts were treated with 1 μM RALF1 for 15 min. Total protein of the protoplasts were extracted and analyzed by SDS‒PAGE and immunoblotting with anti-FLAG, anti-GRP7, anti-GRP7^pS132^ and anti-GFP antibodies.

### The dimerization site mutation FER^P740A^ inhibits FER function

FER plays a pivotal role in the regulation of plant stress responses. To substantiate the impact of mutations at dimerization sites on FER functionality *in vivo*, we conducted hypersensitivity (HR), ROS production and stomatal aperture experiments using the transient expression in plant leaves system to validate these mutant variants (32). Transient transformations of FER or dimerization site mutation variants in *fer-4* leaves were performed. The experimental results indicated a significant hyper-sensitive response exhibited by FER-WT (Figure 6A). Similarly, K565R L742A and D792A exhibited results comparable to FER-WT (Figure 6A). In contrast, mutants at the face-to-face dimerization interface, namely R712A, P740A, and E751A, did not demonstrate notable hyper-sensitive responses, and also in K565R/P740A (Figure 6A). Similar to the results observed in the HR, flg22-induced reactive oxygen species (ROS) production indicated a significant decrease in ROS production in R712A, P740A, E751A and K565R/P740A when compared to the FER-WT (Figure 6B). This suggests that FER may be involved in plant immune responses through allosteric regulation mediated by face-to-face dimerization.

**Fig. 6.**
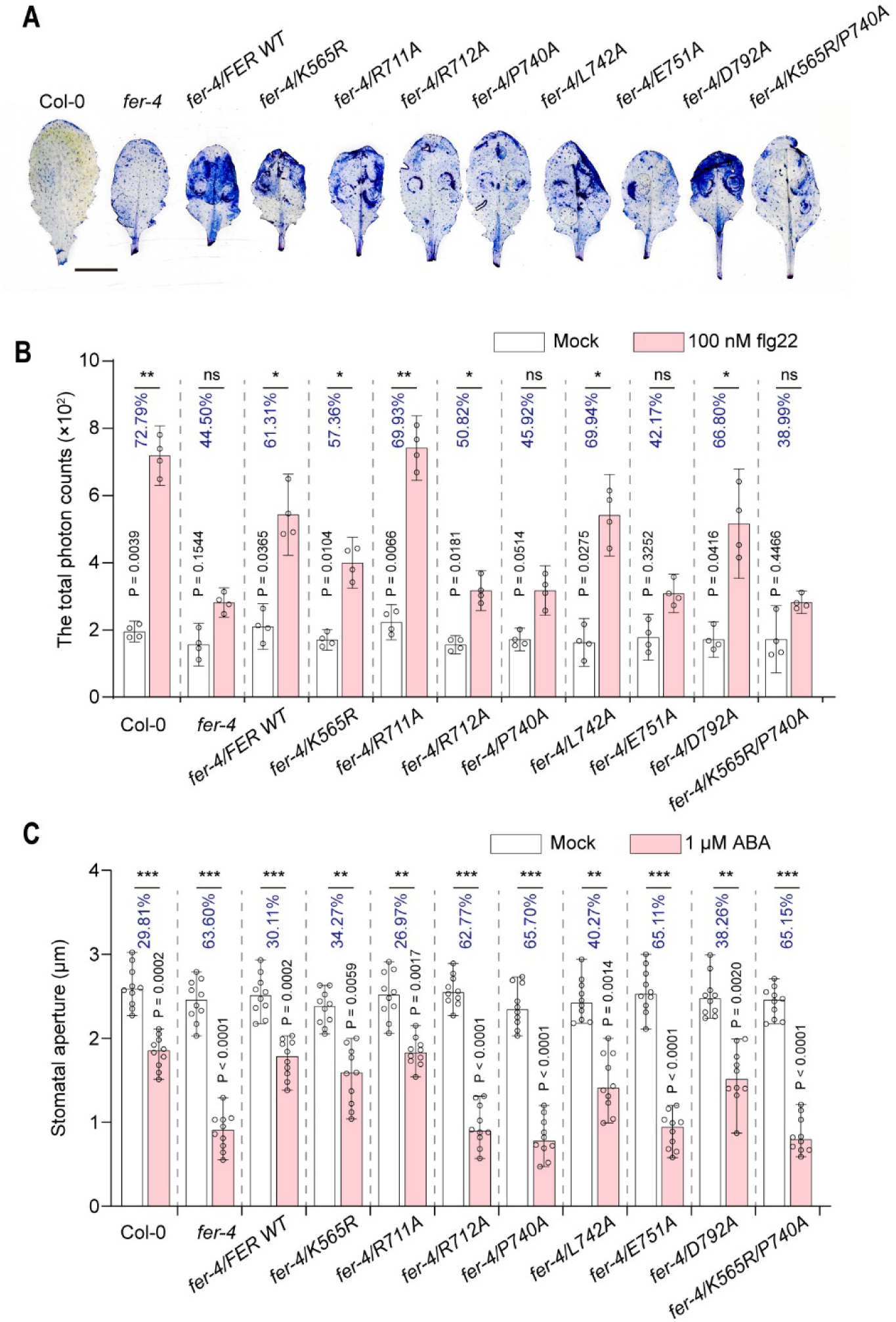
Hypersensitivity (HR) and stomatal aperture of FER variants. (**A**) FER and various mutants fused to C-terminally 4×myc protein were transiently expressed in *fer-4* leaves by Agrobacterium-mediated transient expression (agroinfiltrations). The indicated constructs were agroinfiltrated in *fer-4* leaves at an OD_600_ of 0.5. Pictures of representative leaves exhibit levels of cell death (HR) after injected for 3 days. All assays were performed three times, and a representative photograph is shown. Scale bar, 1 cm. (**B**) FER and various mutants fused to C-terminally 4×myc protein were transiently expressed in *fer-4* leaves by Agrobacterium-mediated transient expression (agroinfiltrations). The indicated constructs were agroinfiltrated in Col-0, *fer-4* and *fer-4* that transiently transformed with FER-WT, K565R, R711A, R712A, P740A, L742A, E751A, D792A and K565R/P740A leaves at an OD_600_ of 0.5. ROS burst measured in leaf discs after stimulation with H_2_O (pH 5.8), 100 nM flg22. The total photon counts of ROS production shown as the mean values recorded over 30 min with SE, n = 4 (one-way ANOVA; *p < 0.05; **p < 0.01; n.s., not significant). All experiments were replicated three times with similar results. (**C**) FER and various mutants fused to C-terminally 4×myc protein were transiently expressed in *fer-4* leaves by Agrobacterium-mediated transient expression (agroinfiltrations). The indicated constructs were agroinfiltrated in *fer-4* leaves at an OD_600_ of 0.5. Stomatal aperture before and after 1 μM ABA treatment. The stomatal aperture shown as the mean values with SE, n = 10 (one-way ANOVA; **p < 0.01; ***p < 0.001). All assays were performed three times.

The previous studies have indicated that, compared to the wild type, ABA can induce stomatal closure in *fer-4* (33). To further validate the impact of R712A, P740A, and E751A on ABA-induced stomatal closure, we utilized the transient expression system in *fer-4* leaves. Col-0 and *fer-4* leaves transiently transformed with FER-WT, K565R, R711A, R712A, P740A, L742A, E751A, D792A and K565R/P740A were exposed to 1 μM ABA treatment for 1 hour after reaching maximum aperture, and stomatal apertures were subsequently measured. The results illustrated that, consistent with *fer-4* sensitivity to ABA-induced stomatal closure, R712A, P740A, E751A and K565R/P740A exhibited a sensitivity in stomatal closure response under ABA induction (Figure 6C). Conversely, transient transformation with FER-WT in *fer-4* leaves restored the ABA-induced stomatal sensitivity phenotype, resembling that of Col-0, and K565R exhibited results comparable to FER-WT (Figure 6C). This suggests that FER may be involved in ABA-induced stomatal closure through allosteric regulation mediated by face-to-face dimerization.

## DISCUSSION

This study provides an in-depth exploration of the allosteric activation mechanism in FER, emphasizing the impactful role of FER^K565R^ and MDS1 on FER and HERK1 kinase activity. These findings settle the debate on FER’s inactivated forms, and reveal a new mechanism for allosteric activation of RLKs via redox signaling, enhancing our understanding of pseudokinases in plants. These findings not only contribute to the understanding of FER regulation but also have broader implications for unraveling the intricacies of RLK signaling pathways.

Previous studies have revealed the roles of RALFs and CrRLK1Ls in modulating ROS production which are essential to plant growth and immunity (29, 34, 35). This study delves into the significance of FER and its homologs in the regulation of ROS signaling. H_2_O_2_ induces significant dimerization in specific CrRLK1Ls family members (Figure 2), and H_2_O_2_ oxidation-induced dimerization enhances the kinase activity of FER and HERK1 (Figure 3A and 3B), highlighting the impact of environmental cues on kinase activity regulation. The discovery of ROS involvement in dimerization reveals its crucial role in allosteric regulation and kinase activity, emphasizing the pivotal position of FER and its family in plant signaling pathways. Furthermore, ROS exhibits a multifaceted role in the signaling of various RLKs, suggesting a potential feedback loop where ROS, once generated and regulated by RLKs, may modulate RLKs to maintain precise signal balance. This intricate interplay underscores the need for future research to explore the regulatory mechanisms governing ROS and RLK interactions within signaling pathways. Understanding these dynamics holds the promise of providing valuable insights into the precise regulation of cellular responses.

In our previous research, we found that unlike the typical mode of protein kinase activation regulation, both FER-WT and FER-K565R exist in a kinase-activated conformation in the unphosphorylated state (26). The ability to maintain this activated conformation independently of phosphorylation regulation suggests its potential significance. This active conformation is expected to play a crucial role in mediating specific functions of the FER, representing a key aspect of FER’s allosteric regulation. Previous genetic studies have shown that kinase-dead FER^K565R^ protein effectively compensated for deficiencies in ovule fertilization, FER-dependent mechanical ion signaling, rosette morphology, and RALF1-mediated stomatal movements. These findings imply that FER’s kinase activity is not essential for these processes. However, dimerization of FER after oxidation is involved in regulating these processes, highlighting the importance of FER’s allosteric activation regulation.

In plants, a substantial number of pseudokinases exist, with the majority of reported and discovered pseudokinases being associated with plant immune regulation. Some non-RD bona fide kinases have been demonstrated to enhance antibacterial immunity in plants, irrespective of phosphotransfer activity (36). Whether these pseudokinases may achieve the output of genetic functions through allosteric regulation, is a topic worthy of further investigation.

In conclusion, this study provides a comprehensive understanding of the allosteric activation of FER and its homologs through the kinase-deficient mutant FER^K565R^. The concentration-dependent enhancement of kinase activity is linked to intermolecular dimerization, especially at the face-to-face interface, and is regulated by ROS signaling. The findings significantly contribute to elucidating the molecular mechanisms governing the allosteric regulation of FER and its homologs, opening avenues for further research in plant signal transduction pathways. Future research focused on unraveling RLK kinase activity regulation (such as allosteric activation) and exploring the impact of specific site variants on genetic and agronomic traits holds tremendous potential for advancing both fundamental plant biology and applied crop improvement efforts.

## METHODS

### Plant materials and growth conditions

All seeds were surface sterilized and stratified at 4°C for 3 days and then grown on half-strength Murashige and Skoog (MS) medium with 0.8% (w/v) sucrose solidified with 1% (w/v) agar (A7002, Sigma-Aldrich) for analysis. Arabidopsis (*Arabidopsis thaliana*) wild-type (Col-0) was the accession used in this study. The *fer-4* mutant and the *pFER::FER-GFP* line in the *fer-4* background were reported previously (35). The *FER^K656R^-GFP* constructs were cloned into the plant expression vectorp1300-LV with *Spe*I and *Pac*I to generate *ACTIN2* promoter-driven constructs. *pFER::FER^K565R^-GFP* plant was crossed to *fer-4*. The *pFER::FER-GFP* face-to-face mutants (R712A, P740A, L742A, E751A and D792A) and *pFER::FER^K565R^-GFP* face-to-face mutants (R712A, P740A, L742A, E751A and D792A) construct were generated in the plant expression vector *p1300-GFP* with *EcoR*I and *BamH*I enzymes, all were crossed to *fer-4*. The *pACTIN2::FER-4×myc*, *pACTIN2::FER^K565R^-4×myc*, and face-to-face mutants (R711A, R712A, P740A, L742A, E751A and D792A) and *pACTIN2::FER^K565R^-GFP* face-to-face mutant P740A construct were generated in the plant expression vector pDT7 with *Spel*I enzyme, positive transformants were selected by resistance to basta and verified by immunoblotting using α-GFP (ABclonal, AE012) antibody. All primers used for plasmid construction and transgenic lines are listed in Table S1.

### Protein expression and purification

FER intracellular domain (FER-CD), FER-CD mutations (K565R, K565R/R712A, K565R/P740A, K565R/E751A, R711A, R712A, P740A, L742A, E751A and D792A, R589A, R591A, Y648A, G652N, A653D, D802A and W805A) and FER homologs intracellular domain (ANJ-CD, ANX1-CD, HERK1-CD, HERK2-CD, MDS1-CD, and MDS2-CD,) were cloned into the pRSF-Duet vector with an N-terminal 6×His tag.

GRP7 was cloned into the pGEX-6P-1 vector. GRP7 and all FER-CD mutations or FER homologs were co-expressed with λ-phosphatase (λ-PP), which was cloned into the pGEX-6P-1 vector and pRSF-Duet vector, respectively. All proteins were expressed in the same *E. coli* strain, BL21(DE3). Protein expression was induced at 18℃ for 18 h with 1 mM isopropyl-β-D-thiogalactopyranoside (IPTG) after the OD_600_ reached 0.6. The cells were harvested by centrifugation at 5000 rpm for 15 min, resuspended in phosphate-buffered saline (PBS, pH 7.5) containing 10 mM imidazole, 1 mM DTT and crushed by using a low-temperature ultrahigh-pressure cell disrupter (JNBIO, Guangzhou, China). The soluble proteins were separated from cell debris by high-speed centrifugation (12,000 rpm for 1 h). 6×His-tagged proteins were loaded onto a Ni^2+^ affinity column (Smart Life Sciences, SA004250) and washed with 100 ml of washing buffer (PBS containing 20-100 mM imidazole). The target protein was eluted from the column with PBS containing 300 mM imidazole. GRP7 proteins were hung on GST beads (Smart Life Sciences, SA008100), and washed with 60 ml of washing buffer (PBS 7.5). GRP7 protein was digested overnight with PreScission Protease (YBscience, YB11401) and eluted with PBS. All proteins were exchanged into HEPES buffer (50 mM HEPES, 10% Glycerol; pH 7.5) by using centrifugal concentrators (30 kDa, Millipore, UFC901096).

### *In vitro* kinase assay

For *in vitro* kinase assays, all samples were incubated with 1 mM ATP and 10 mM Mg^2+^ at 25°C for 40 min or for the specified time gradient. The samples were mixed with 4×loading buffer (v/v), and boiled at 98°C for 5 min. All samples were separated by SDS-PAGE and either stained with Coomassie blue G250 or transferred to a nitrocellulose membrane (PALL, P-N66485) for protein immunoblotting using anti-phospho-Serine/Threonine (ABclonal, AP0893) antibody.

### *In vivo* kinase assay

For the *in vivo* kinase assay, anti-GRP7 and anti-GRP7^pS132^ antibodies were generated by ABclonal (Wuhan, China) (37). Protoplasts were prepared from 7-day-old Col-0, *fer-4* or *Ubi::FER-FLAG* seedlings. About 40 μg of DNA plasmids were transfected into protoplasts. The transfected protoplasts were incubated in the dark at 23°C for 18 h to allow protein production. RALF1 was synthesized by Guoping Pharmaceutical Co., Ltd., and dissolved in water. For RALF1-induced phosphorylation of FER, the transfected protoplasts were treated with 1 μM RALF1 for 15 min. Total protein of the protoplasts were extracted and analyzed by SDS‒PAGE and immunoblotting with anti-GRP7, anti-GRP7^pS132^, anti-FLAG (ABclonal, AE005) and anti-GFP antibodies.

### BiFC assays

For the BiFC assay, FER and HERK1-N-terminal Venus (nVenus) were amplified by PCR and cloned into the pE3308 plasmids, and FER^K565R^ and MDS1-C-terminal (cCFP) were amplified by PCR and cloned into the pE3449 plasmids, respectively. Protoplasts were isolated from well-expanded rosette leaves of 5-weekold Arabidopsis plants through cellulase and macerozyme digestion. Then protoplasts were co-transfected with the nVenus and cCFP constructs, followed by incubation in the dark at 23℃ for 18 h. Fluorescence was monitored with a confocal microscope using an excitation wavelength of 488 nm for GFP.

### Split-luciferase complementation imaging (LCI) assay

Each of the constructs containing full-length cDNAs were cloned with split-LUC at their C-termini of the *pCAMBIA1300-GW-nLUC* or *pCAMBIA1300-GW-cLUC* vector and transformed into the GV3101 by GenePluserTM electro-transformation (BioRad). When the OD600 reached 1.0, the cultures were resuspended with infiltration buffer [10 mM MES (pH = 5.7), 10 mM MgCl_2_, 100 μM 4’-Hydroxy-3’,5’-20 dimethoxyacetophenone] and incubated for 2 h. GV3101 carrying the indicated constructs (OD600 = 0.6) was infiltrated into 4-week-old leaves of *N. benthamiana*. The infiltrated leaves were incubated at 28°C for 2 days before LUC activity measurement. The luminescence was monitored and captured using a low light imaging CCD camera (Photek; Photek, Ltd.).

### Agrobacterium-mediated transient transformation

Agrobacterium-mediated transient transformation was performed as described previously (32). The agrobacteria were grown on YEB-induced medium plates at 28℃ for 24–36 h. The cells were scraped and resuspended in 500 μl washing solution (10 mM MgCl_2_, 100 mM acetosyringone). The agrobacteria were finally diluted to an OD600 of 0.5 in infiltration solution (1/4 MS [pH = 6.0], 1% sucrose, 100 μM acetosyringone, 0.005% [v/v, 50 μl/l] Silwet L-77). The agrobacteria were infiltrated into Arabidopsis leaves using a 1-ml plastic syringe, kept in the light to dry the leaves (1 h), and then subsequently kept in the dark for 24 h at room temperature. The transformed plants were then transferred back to the greenhouse for another 2–3 days before sample microscopy. The efficiency was calculated by observing all cells under a confocal microscope in the same focal plane with four biological replicates.

### Stomatal aperture and ROS assay

Stomatal aperture assays was performed as described previously (38), leaves were floated on stomatal opening buffer (5 mM MES, 5 mM KCl, 50 μM CaCl_2_, pH 5.6) for 3 h in the light (light intensity about 150 μmol m^-2^ s^-1^) to initially open stomata, then treated with 1 μM ABA for 1 h. Images of stomatal apertures were taken by Olympus IX73 inverted microscope and measured in each image using ImageJ.

The flg22-induced ROS burst assay, was performed as described previously (39). Four-week-old leaves were cut into 4×4 mm pieces that were transferred to 96-well plates containing 100 μl of sterile water and then covered and incubated in the dark overnight. After 12 h, the water was replaced with 100 μl of a solution containing 20 μM luminol L-012 (Sigma-Aldrich), 1 μg/ml horseradish peroxidase (Sigma-Aldrich), 100 nM flg22 (PhytoTech, Lenexa, KS, USA) or elution buffer. Luminescence was measured using a Fluoroskan Ascent FL microplate fluorometer and luminometer (Thermo Scientific). ROS production was reported as either the increase in photon counts or the sum of total photon counts.

### Sequence alignment

The amino acid sequences of FER subfamily proteins from Arabidopsis thaliana were obtained from The Arabidopsis Information Resource (TAIR) site. The sequences were subjected to multiple sequence alignment (MSA) using the T-Coffee server (40), and the MSA results were visualized using ESPript version 3.0 (41).

### Quantification and statistical analysis

For quantifying the nucleus-localized fluorescence intensity of MAMP markers, confocal images were analyzed with the Fiji package (http://fiji.sc/Fiji). Contrast and brightness were adjusted in the same manner for all images. All statistical analyses were done with Graphpad Prism 8.0 software. One-way analysis of variance (ANOVA) was performed, and Tukey’s test was subsequently used as a multiple comparison procedure. Details about the statistical approaches used can be found in the figure legends. The data are presented as means ± standard deviation (SD), and ‘‘n’’ represents the number of plant roots.

## Acknowledgments

We thank Dr. Alice Cheung for providing plant materials

## Funding

This work was supported by grants from National Natural Science Foundation of China (NSFC-32000916), Natural Science Foundation of Hunan Province (2021JJ40050), China Postdoctoral Science Foundation (2019M662764)

## Authors’ contributions

F. Y. and J. C. conceived the project and designed research; J. C, D. C, Y-X. Z, L. C performed research; J. C. and F. Y. analyzed data and wrote the paper; all authors reviewed and approved the manuscript for publication.

## Competing interests

Authors declare that they have no competing interests.

## Data and materials availability

All data are available in the manuscript or the supplementary materials.

## Supplementary Materials

Materials and Methods

Figs. S1 to S4

Table S1

References

**Fig. S1.**
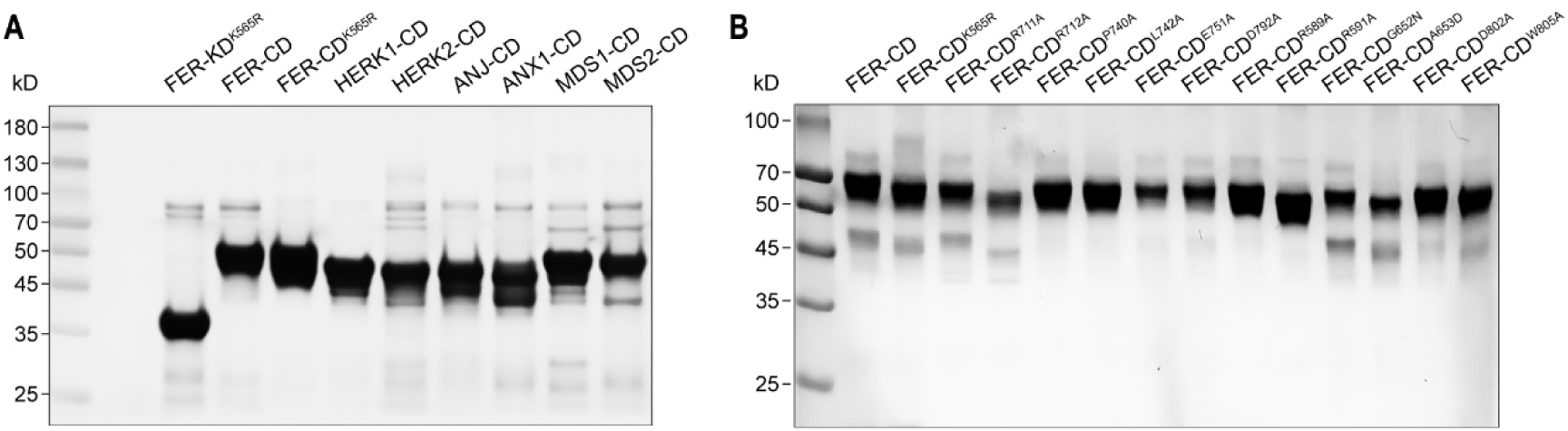
SDS–PAGE analysis of purified FER-CD, FER-CD mutants and FER subfamily proteins. (**A**) FER-CD and FER subfamily proteins in cytoplasmic kinase domain (CD) were cloned into the pRSF-Duet vector with an N-terminal 6×His tag, and were co-expressed with λ-phosphatase (λ-PP), which was cloned into the pGEX-6P-1 vector. All proteins were expressed in the same *E. coli* strain, BL21(DE3) and exchanged into HEPES buffer (50 mM HEPES, 10% Glycerol; pH 7.5) by using centrifugal concentrators (30 kDa, Millipore, UFC901096). (**B**) FER-CD, face-to-face, and back-to-back dimer interface mutants were cloned into the pRSF-Duet vector with an N-terminal 6×His-FRB tag (FER-CD) or FKBP tag (FER-CD mutants), and were co-expressed with λ-phosphatase (λ-PP), which was cloned into the pGEX-6P-1 vector. All proteins were expressed in the same *E. coli* strain and exchanged into HEPES buffer by using centrifugal concentrators.

**Fig. S2.**
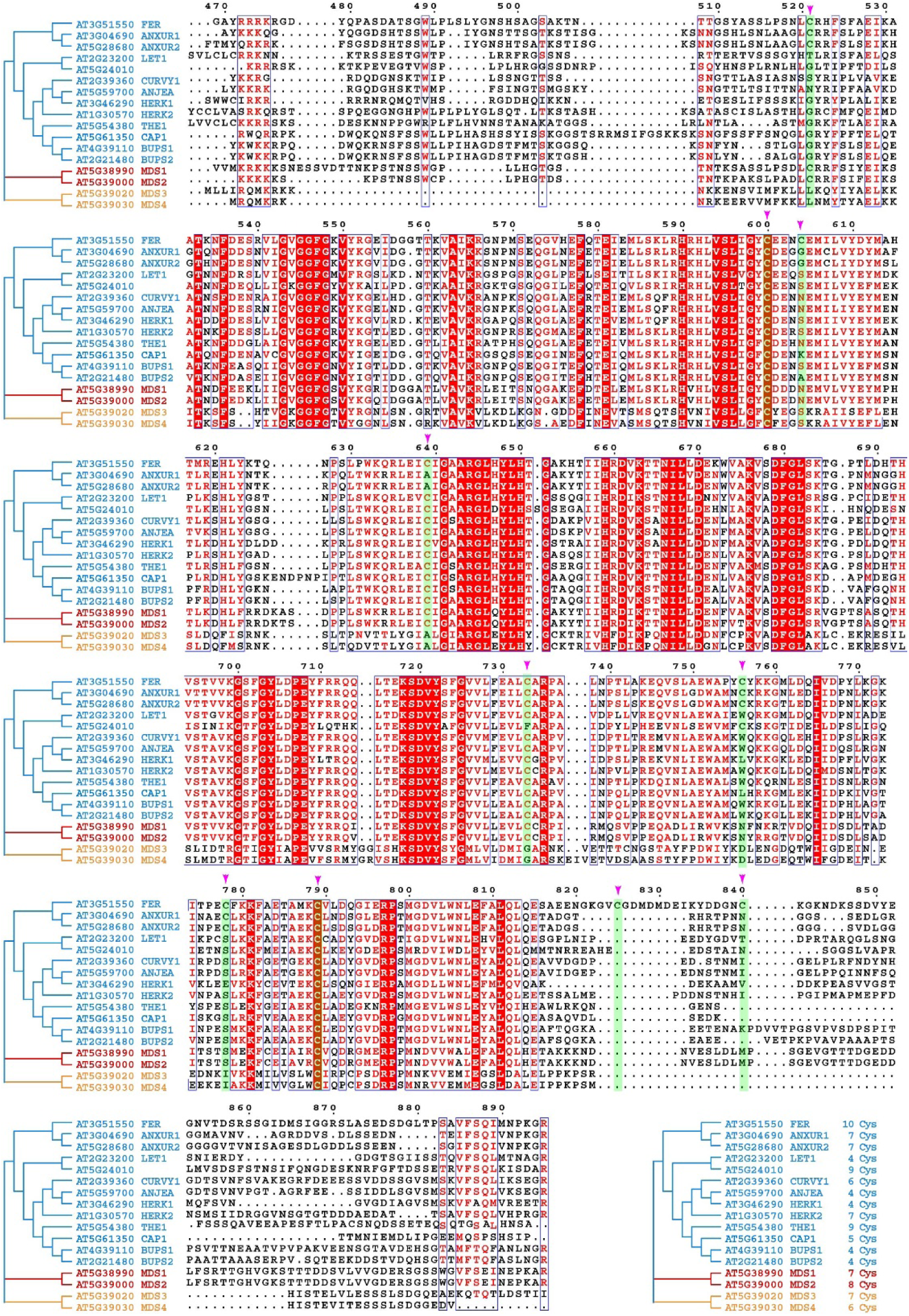
Sequence alignment of FER-CD with sixteen representative FER subfamily proteins in CD. The 10 cysteine residues distribution in FER-CD are shown with pink arrow. The number and distribution of cysteine residues of FER-CD and homologous family members are very different. FER-CD contains 10 cysteine residues, HERK1-CD contains 4 cysteine residues, HERK2-CD contains 7 cysteine residues, ANJ-CD contains 4 cysteine residues, ANX1-CD contains 7 cysteine residues, MDS1-CD contains 7 cysteine residues, and MDS2-CD contains 8 cysteine residues.

**Fig. S3.**
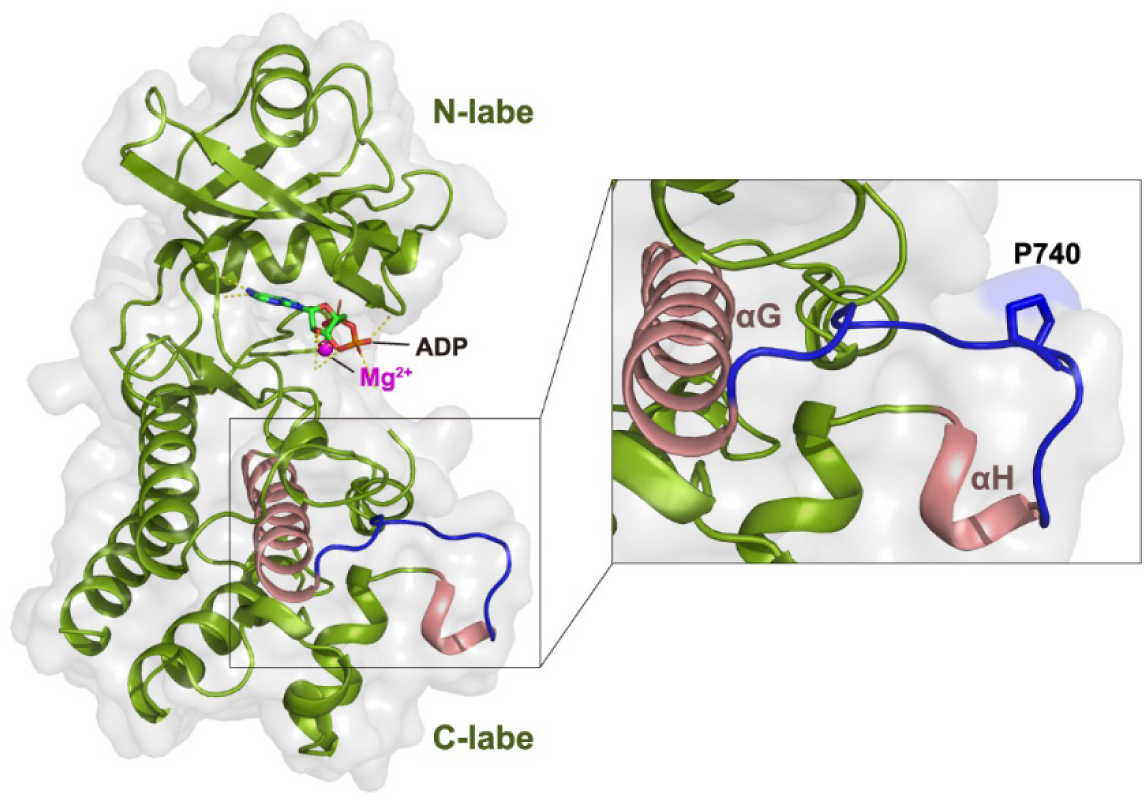
Detailed presented of the P740 residue within the *α*G-*α*H loop. The structure of FER-KD, as shown in a cartoon representation. ADP is shown as colored sticks, and Mg^2+^ ions are indicated by purple spheres. P740 residue is presented as blue sticks. αG and αH presented as pink helix, αG-αH loop presented as blue loop.

**Fig. S4.**
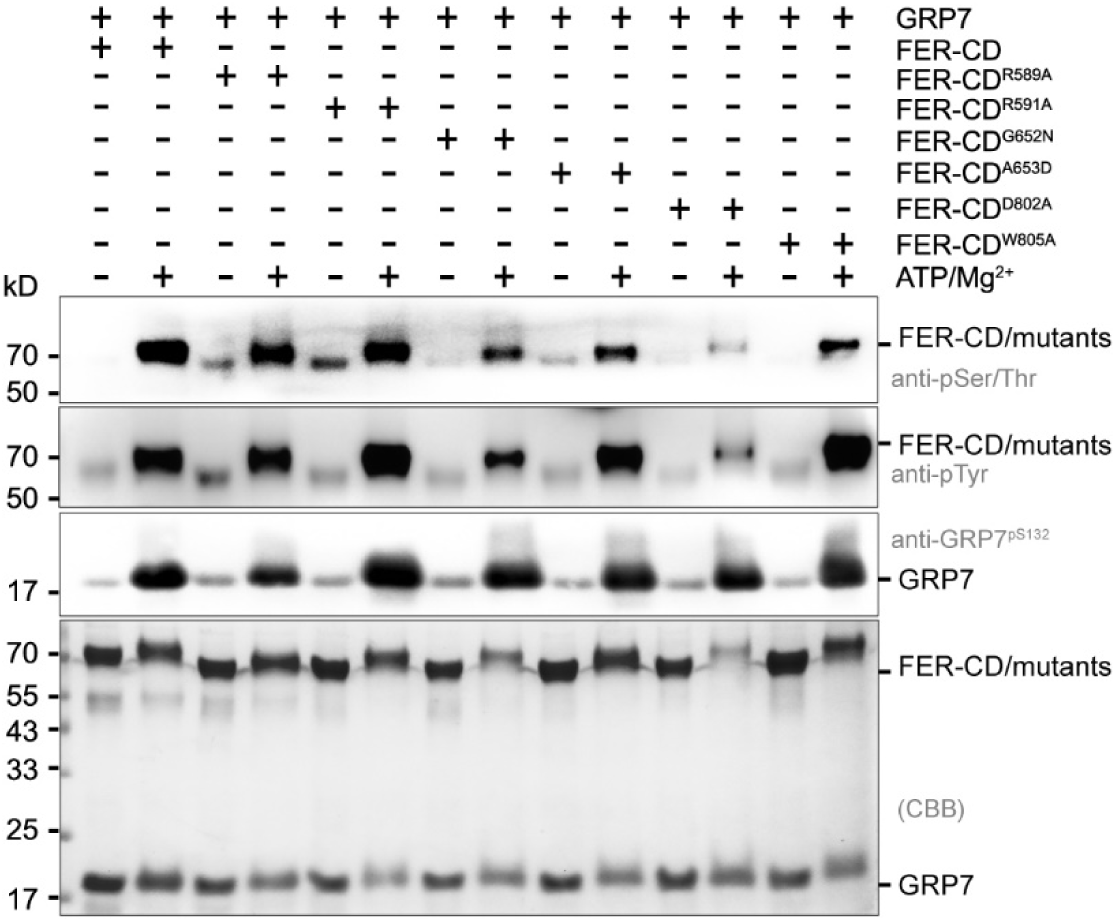
The back-to-back dimerization interface sites of FER-CD are not crucial for FER kinase activity. Western blot analysis of autophosphorylation and transphosphorylation of back-to-back dimerization interface sites of FER-CD. Samples were incubated with 1 mM ATP and 10 mM Mg^2+^ at 25°C for 30 min.

## Notes

### Competing Interest Statement

The authors have declared no competing interest.

